# Are adaptive chemotherapy schedules robust? A three-strategy stochastic evolutionary game theory model

**DOI:** 10.1101/2021.02.28.433232

**Authors:** R. Dua, Y. Ma, P.K. Newton

**Affiliations:** Department of Mathematics, University of Southern California, Los Angeles CA 90089-1191; Department of Physics & Astronomy, University of Southern California, Los Angeles CA 90089-1191; Department of Aerospace & Mechanical Engineering, Mathematics, and The Ellison Institute, University of Southern California, Los Angeles CA 90089-1191

## Abstract

We investigate the robustness of adaptive chemotherapy schedules over repeated cycles and a wide range of tumor sizes. We introduce a non-stationary stochastic three-component fitness-dependent Moran process to quantify the variance of the response to treatment associated with multidrug adaptive schedules that are designed to mitigate chemotherapeutic resistance in an idealized (well-mixed) setting. The finite cell (*N* tumor cells) stochastic process consists of populations of chemosensitive cells, chemoresistant cells to drug 1, and chemoresistant cells to drug 2, and the drug interactions can be synergistic, additive, or antagonistic. First, the adaptive chemoschedule is determined by using the *N* → ∞ limit of the finite-cell process (i.e. the adjusted replicator equations) which is constructed by finding closed treatment response loops (which we call evolutionary cycles) in the three component phase-space. The schedules that give rise to these cycles are designed to manage chemoresistance by avoiding competitive release of the resistant cell populations. To address the question of how these cycles are likely to perform in practice over large patient populations with tumors across a range of sizes, we then consider the statistical variances associated with the approximate stochastic cycles for finite *N*, repeating the idealized adaptive schedule over multiple periods. For finite cell populations, the error distributions remain approximately multi-Gaussian in the principal component coordinates through the first three cycles, with variances increasing exponentially with each cycle. As the number of cycles increases, the multi-Gaussian nature of the distribution breaks down due to the fact that one of the three subpopulations typically saturates the tumor (competitive release) resulting in treatment failure. This suggests that to design an effective and repeatable adaptive chemoschedule in practice will require a highly accurate tumor model and accurate measurements of the subpopulation frequencies or the errors will quickly (exponentially) degrade its effectiveness, particularly when the drug interactions are synergistic. Possible ways to extend the efficacy of the stochastic cycles in light of the computational simulations are discussed.

**Prepared for Special Issue:** *Understanding the Evolutionary Dynamics and Ecology of Cancer Treatment Resistance*, Ed. D. Basanta, Cancers (2021)

## I. INTRODUCTION

The design of adaptive chemotherapy schedules [1], motivated and aided by mathematical/computational models [2], is a rapidly developing field that has great potential for mitigating chemoresistance in tumors [3]. The advantage of an adaptive schedule over a more standard pre-determined schedule, such as a maximum tolerated dose (MTD) schedule, or a low-dose metronomic (LDM) schedule [4, 5], is that adaptive schedules are able to evolve along with the tumor [6]. But as has been observed [7], the efficacy of sequential adaptive cycles depends crucially on the ability to monitor and track the heterogeneous subpopulations of cells that comprise the tumor [8], or at least an accurate surrogate biomarker [9, 10], which can be challenging. Additionally, the fact that any complex finite population of cells will evolve stochastically [11–13] makes it particularly important to assess the effectiveness of adaptive schedules under a range of diverse conditions, such as different patient populations and tumor sizes and over many therapy cycles. We present a mathematical model that addresses these issues.

Standard pre-scheduled chemotherapy dose-delivery schedules suffer from the common occurence of chemoresistance of the tumor [14], leading to treatment failure [15] after multiple cycles. To overcome this, adaptive schedules are designed with the goal of managing resistance [1]. Adaptive therapies typically leverage tumor heterogeneity [8] by exploiting evolution via competition of the tumor subpopulations [16] to *steer evolution* and designing an advantageous chemoschedule [17–19] that delays chemoresistance by maintaining a sufficient fraction of the sensitive population in order to supress the resistant population, a strategy used in the context of avoiding antibiotic resistence [20] as well.

Aside from the schedule (dose and timing), the use of two or more drugs is helpful in mitigating resistance, if combined optimally [2, 21]. A key challenge associated with dose scheduling and the optimal design of multidrug combinations is that both the drug schedules and the multidrug mixture rely on an accurate dynamical assesment of the relative balance and mixture of the evolving cell types in the tumor which is hard to obtain in clinical practice [7]. Quantitative assessments are, generally speaking, much easier to obtain in a mathematical model. But even within the framework of a mathematical model, how exactly to quantify the statistical effectiveness of the schedule and multidrug mixture when the schedule is adaptive (i.e. is not fixed and repeatable) is not at all straightforward. We address this important issue of uncertainty quantification and robustness [22–24] of tumor response to adaptive therapy schedules and synergystic vs. antagonistic multidrug interactions [25, 26] by using a stochastic finite-cell fitness-dependent Moran process evolutionary game theory tumor model with an adaptive schedule designed from the deterministic adjusted replicator dynamical system [27], which is the large cell limit (*N*→ ∞) of the finite-cell stochastic process [28, 29]. We describe the model as well as the connections between the finite cell stochastic model and infinite cell deterministic model in the next section.

## II. THREE COMPONENT TUMOR CELL KINETICS MODEL

### A. Three-component adjusted replicator dynamics

The deterministic model we use to determine the sub-population frequencies (*x*_1_, *x*_2_, *x*_3_) ≡ (*S, R*_1_, *R*_2_) and the dosing schedules (*C*_1_ (*t*), *C*_2_ (*t*)) to determine the adaptive therapy are the adjusted replicator equations:

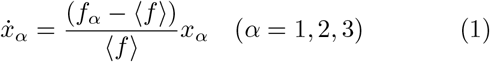

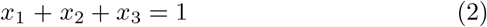

Here, *f*_*α*_ denotes the fitness of subpopulation *α*:

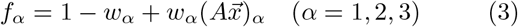

While 〈*f*〉 denotes the average fitness of the entire population (tumor):

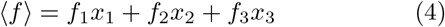

Thus, the growth or decay of population *α* in eqn (1) is determined by the deviation of the fitness of population *α* from the average, normalized by the average fitness of the tumor.

The relative fitness of the three subpopulations are controlled by the selection pressure parameters *w*_*α*_:

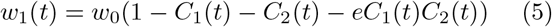

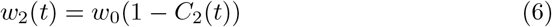

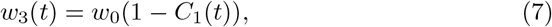

Where 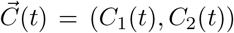) is the chemotherapy delivery function, with *C*_1_ controlling drug 1, *C*_2_ controlling drug 2, and *e* is a parameter that determines whether the two drugs act synergistically (*e >* 0), antagonistically (*e <* 0), or additively (*e* = 0). We take *w*_0_ = 0.1 which sets the timescale in our simulations. Notice that when drug 1 is applied (*C*_1_(*t*) *>* 0), it acts to reduce the selection pressure parameters *w*1, *w*3, but leaves *w*2 (the parameter controlling the *R*1 population) unchanged, so the *R*1 population is resistant to drug 1. When drug 2 is applied (*C*_2_ (*t*) *>* 0), it acts to reduce the selection pressure parameters *w*_1_, *w*_2_, leaving *w*_3_ unchanged (the parameter controlling the *R*_2_ population), so the *R*_1_ population is resistant to drug 2. Our model steers evolution by exploiting the effect that chemotherapy has on the selection pressure applied to different subpopulations. The total dose delivered over time-period *t* is denoted 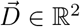:

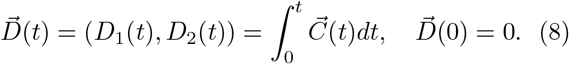

Then:

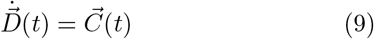

and:

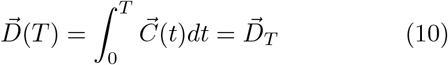

Thus,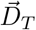 denotes the total dose delivered in fixed time *T*. The system (1) is a nonlinear non-constant coefficient dynamical system governing the frequency distributions.

To fix the evolutionary game being played by the population of cells, we consider the 3 × 3 payoff matrix *A*:

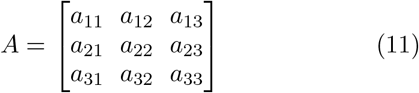

In order to model tumor kinetics, we take our payoff matrix to be of Prisoner’s Dilemma (PD) type [30], with conditions:

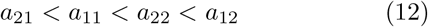

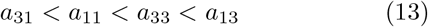

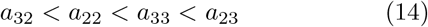

For definiteness, we choose the specific values:

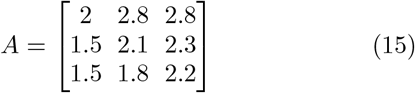

The PD payoff matrix is what makes the model useful as a cancer model, where the sensitive cell population *S* plays the role of defectors, and the two resistant cell populations *R*_1_ ane *R*_2_ play the role of cooperators, with lower (but unequal) fitness. The values we choose build in a cost to resistance [31], which allows the sensitive cell population to saturate the tumor in the absence of chemotherapy (*C*_1_ (*t*), *C*_2_ (*t*)) = (0, 0). Our chemotherapy functions are constrained so that 0 ≤ *C*_1_(*t*) ≤ 1, 0 ≤ *C*_2_(*t*) ≤ 1, 0 ≤ *C*_1_ (*t*) + *C*_2_ (*t*) ≤ 1, and from eqns (5)-(7) reduce the fitness of the relevant subpopulations by altering the relative selection pressures.

As a final equation comprising our model, we have a tumor growth equation, where *V* (*t*) represents the total tumor volume:

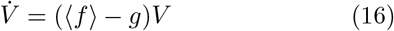

Here, *g* denotes a positive constant we take as generally representing the fitness of the average microenvironment surrounding the tumor. Hence, the growth rate (or decay) of the tumor, (〈*f*〉 − *g*), is given by the deviation of the average tumor fitness from the average fitness of the local microenvironment, as represented by the gray region surrounding the tumor cells in figure 1.

**FIG. 1.**
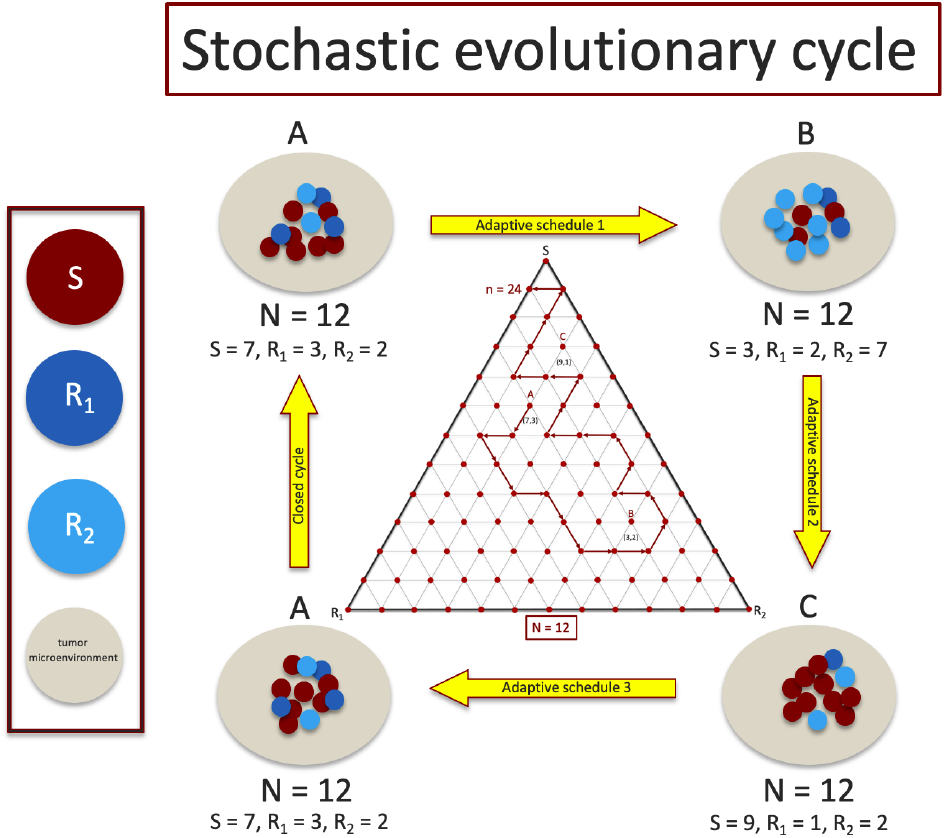
A sequence of adaptive chemoschedules that lock the tumor in a closed evolutionary cycle is difficult to achieve for finite cell (shown for *N* = 12) populations since subpopulation frequencies fluctuate stochastically and are difficult to measure with precision. Middle inset shows discrete trilinear plot of a stochastic realization in (*S, R*_1_, *R*_2_) plane, with *n* = 24 steps, starting at point *A*, using the chemoschedule designed from the deterministic (*N* → ∞) model.

### B Three-component discrete stochastic process

Our finite cell model is a three component stochastic birth-death process [32, 33], with frequency-dependent fitness governed by a payoff matrix, and population size *N* comprised of a fluctuating mixture of cells of type *S* (sensitive), cells of type *R*_1_ (resistant to drug 1), and cells of type *R*_2_ (resistant to drug 2), with *S*+*R*_1_+*R*_2_ = *N*. At every step *n* in the process, a cell is randomly selected for reproduction with probability proportional to its fitness. The offspring produced replaces a randomly chosen cell in the population. The fitness of cells in the subpopulations *S, R*_1_, and *R*_2_ are given respectively as:

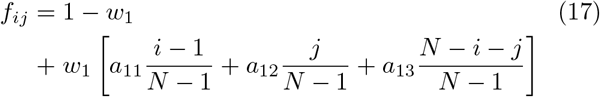

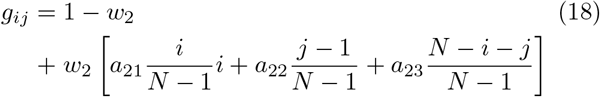

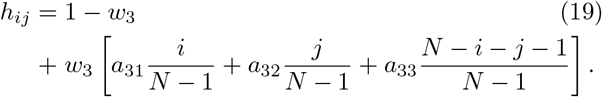

Here, *i* denotes the number of cells of type *S, j* denotes the number of cells of type *R*_1_, and *N*—*i*−*j* denotes the number of cells of type *R*_2_, with 0 ≤ *i* ≤ *N*; 0 ≤ *j* ≤ *N, i* + *j* ≤ *N*. The state of the system is given by (*i, j*) (visualized as a grid point on a triangular grid as in figure 1) with probability of residing in that state after *n* steps given by 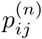.

As in the continuous model, the parameter *w*_*α*_∈ [0, 1] (*α* = 1, 2, 3) controls the selection pressure and allows us to independently adjust the fitness landscape. At the two extremes, when *wα* = 0, the payoff matrix makes no contribution to fitness, so only random drift governs the dynamics. At the other extreme, when _*wα*_ = 1, the payoff matrix makes a large contribution to the fitness, with selection pressure driving the dynamics. We define the average tumor fitness, 〈*f*〉_*N*_, as the discrete analogue of (4):

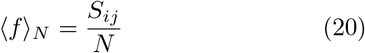

where:

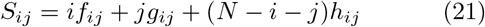

The transition probabilities at each step of the birth-death (Markov) process are written as:

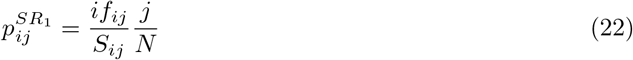

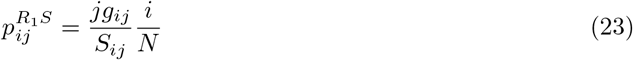

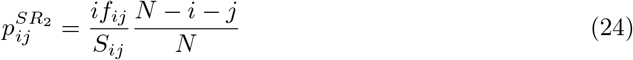

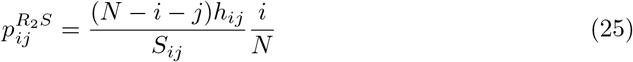

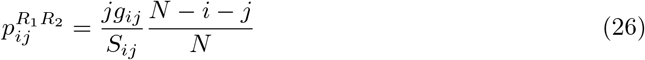

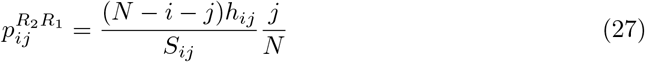

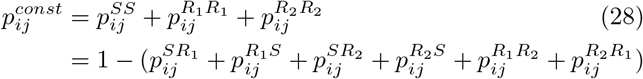

To implement chemotherapy in the discrete Markov process, we discretize time 0 ≤ *t* ≤ *T* in (8)-(10) so that at each time-step *n* in the process (with *n* = *tN*), we enforce the time-dependent chemotherapy schedules (5)-(10) by adjusting the selection pressures where:

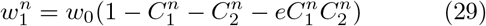

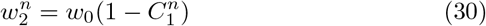

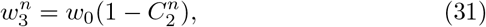

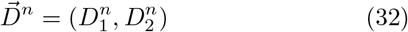

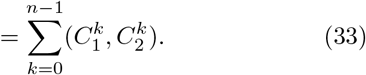

This renders the Markov process non-stationary [34]. The state-vector 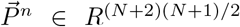 represents the discrete probability distribution at each triangular grid point 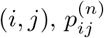 (the finite analogue of the Master equation and Fokker-Planck formulation in [29]):

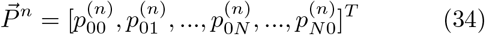

along with a 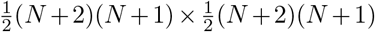 stochastic (non-stationary) transition matrix *M* ≡ {*p*^*ij*^} (whose entries are eqns (22)-(31)) with rows adding to 1 driving the dynamical system:

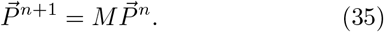

We initiate the process with an initial discrete distribution 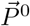 This framework allows us to carry out Monte Carlo simulations to generate probability distributions (using eqns (22) - (27) more practically) of the three cell types for any initial distribution.

### C. Continuous limit *N* → ∞

The deterministic and stochastic models are connected by the limit *N* → ∞ in which the discrete finite cell frequency-dependent model (35) converges to the adjusted replicator system (1). This connection was first made explicit in [28, 29] with an earlier approach in [35]. These papers accomplished two things. First, the authors of [28] showed that different microscopic stochastic processes for two-strategy games lead to either the replicator system or the adjusted replicator system in the limit *N*→ ∞. They also derived an explicit mean-field Fokker-Planck equation describing the probability density function for the different subpopulations. This work was extended in [29] to an arbitrary number of strategies (subpopulations), with the possible inclusion of mutations leading to the adjusted replicator dynamics as *N*→ ∞. Our method of quantifying the uncertainty associated with adaptive chemotherapy schedules exploits this stochastic-deterministic connection. The deterministic adjusted replicator equation (1) allows us to design a schedule giving rise to the closed evolutionary cycle, while the stochastic fitness-dependent three-strategy Moran process allows us to quantify the variance associate with repeated cycling of the schedule for finite *N*.

## III. RESULTS

In figure 2(a) we show the closed deterministic evolutionary cycle *ABCA* in the (*S, R*_1_, *R*_2_) tri-linear plane, along with two stochastic realizations starting at point *A* but not closing the loop. Figure 2(b) depicts the chemoschedule that produces the closed *ABCA* cycle, which we use as our adaptive chemo-schedule for the finite *N* model, repeating it in a series of cycles. In figure 3(a) we show the spread of the endpoints in the (*S, R*_1_, *R*_2_) tri-linear plane after one evolutionary cycle, for the case *e* = 0 when the two drugs act additively on the population. The cycle is initiated at (*S, R*_1_, *R*_2_) = (0.8, 0.1, 0.1) (point marked *A* in figure 2), for 10, 000 trials, with *N* = 100, 000 cells. Also shown are the two orthogonal principal components associated with the spread of the data, which we then map, in figure 3(b) to the two principal axes. The kernel density estimates (KDE) for the distributions are shown in figures 3(c),(d). The distributions, after one loop, closely resemble multivariate Gaussian distributions in the principal components.

**FIG. 2.**
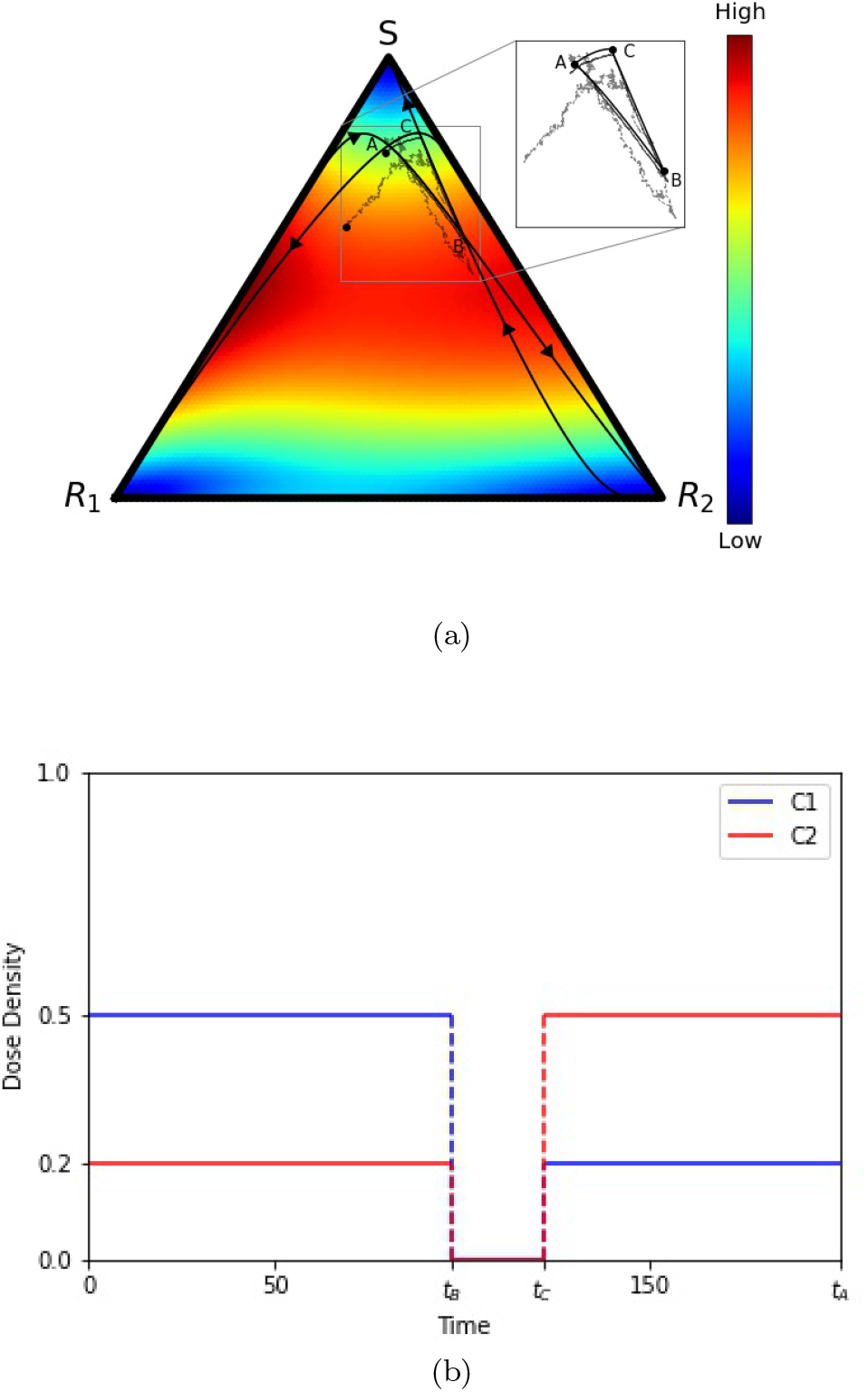
(a) Trilinear (*S, R*_1_, *R*_2_) coordinate representation of a deterministic evoutionary cycle *ABCA*, from the adjusted replicator equation, and two stochastic realizations (from the Moran process) with *N* = 10, 000 and *N* = 1, 000, 000 cells. Inset shows that the stochastic paths are not closed. Background colors show velocity field, hence instantaneous speed of convergence, from the adjusted replicator system.; (b) Adaptive chemotherapy schedule with drugs 1 and 2 for closed cycle *ABCA*.

**FIG. 3.**
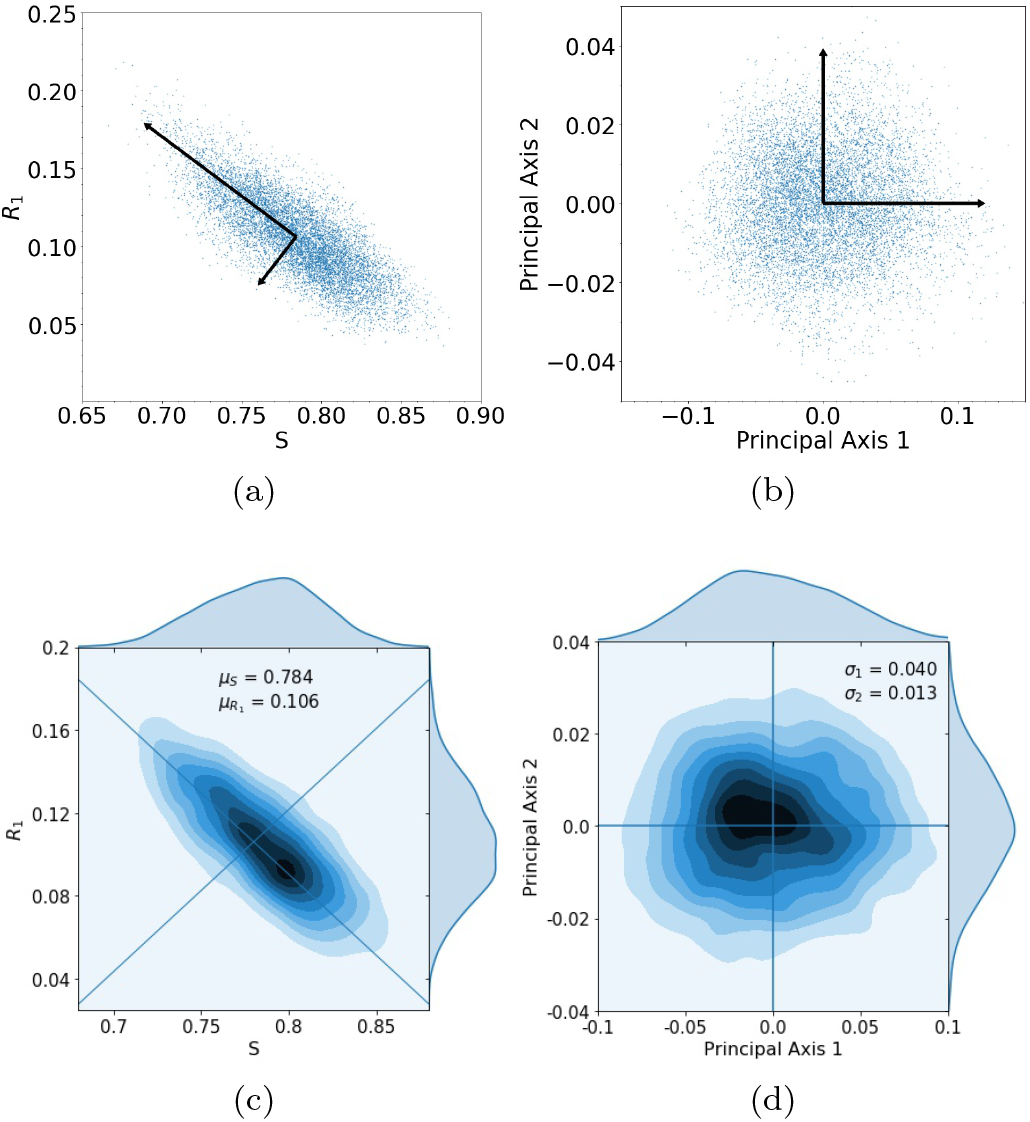
(a) Spread of 10,000 trials starting at initial condition *S* = 0.8, *R*_1_ = 0.1, and *R*_2_ = 0.1 with principal axis after 1 loop.; (b) Same spread of trials from (a) plotted with principal component axes.; (c) Kernel density estimation (KDE) for trials in (a), with darker areas indicating a higher concentration of data, with means *µ*_*S*_ = 0.784, *µ*_*R*1_ = 0.106.; (d) The spread of trials closely resembles a multivariate Gaussian distribution composed of the principal components, with *σ*_1_ = 0.040, *σ*_2_ = 0.013.

After more loops of the cycle, the multivariate Gaussian distribution starts to break down, as shown clearly in figure 4 showing the KDE after 2 loops (4(a)), 3 loops (4(b)), 4 loops (4(c)), and 5 loops (4(d)). After the third loop, the distribution no longer approximates a multi-Gaussian.

**FIG. 4.**
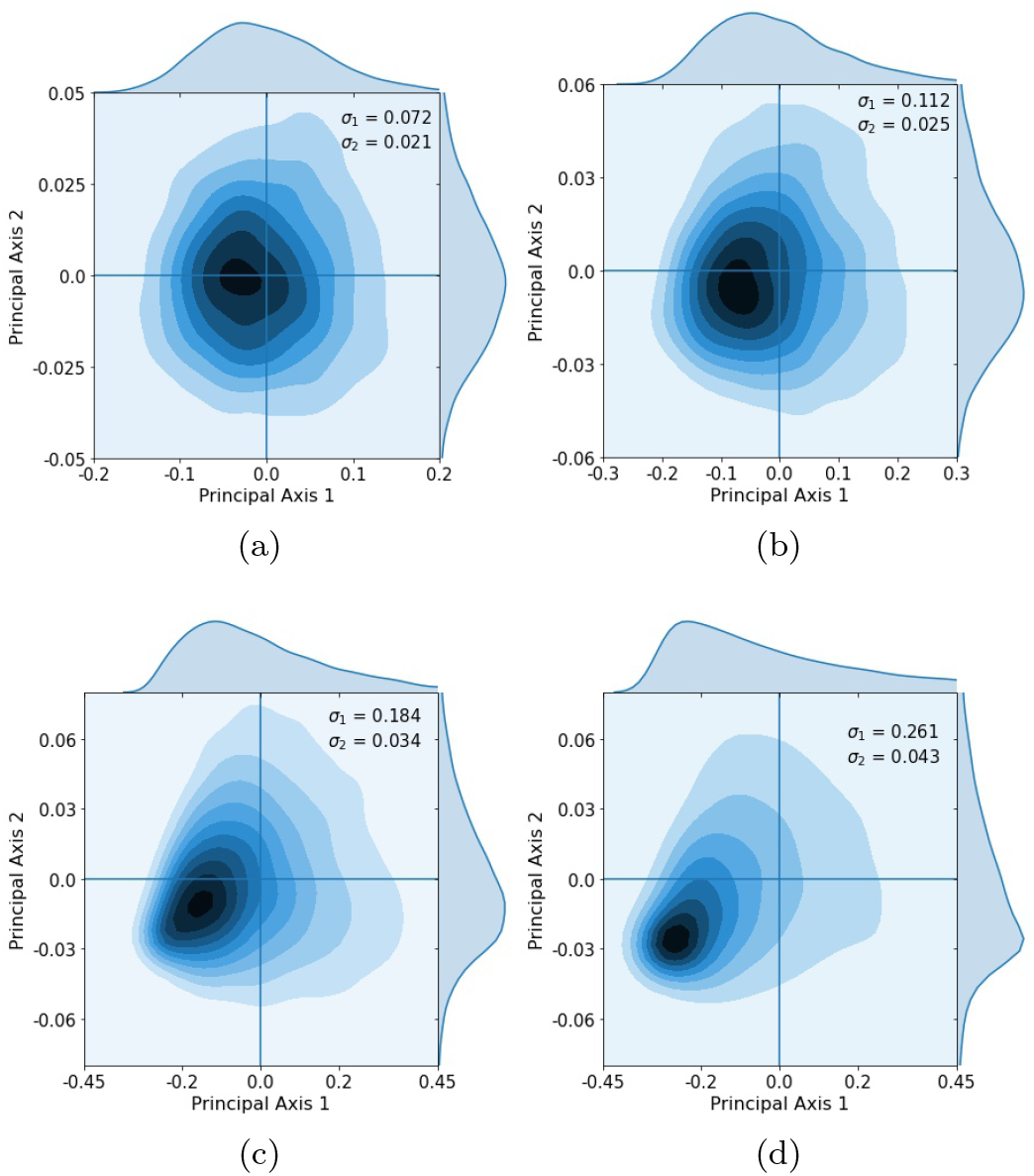
Kernel density estimation showing distribution after successive loops. Gaussian spread starts to break down after loops 3 and 4 as some tumors began to saturate. (a) Loop 2, *σ*_1_ = 0.072, *σ*_2_ = 0.021.; b) Loop 3, *σ*_1_ = 0.112, *σ*_2_ = 0.025,; (c) Loop 4, *σ*_1_ = 0.184, *σ*_2_ = 0.034.; (d) Loop 5, *σ*_1_ = 0.261, *σ*_2_ = 0.043.

To examine why the multi-Gaussian nature of the distribution breaks down, we show in figure 5 histogrammed distributions of the three subpopulation frequencies. By the third loop, the distributions start to impinge on 0 (one of the sides of the tri-linear plane) or 1 (one of the corners of the tri-linear plane), indicating that a subpopulation either vanishes or saturates, distorting the multi-Gaussian spread. By the fourth and fifth loops, the distributions start to converge at these endpoints and the distribution no longer resembles a multi-Gaussian.

**FIG. 5.**
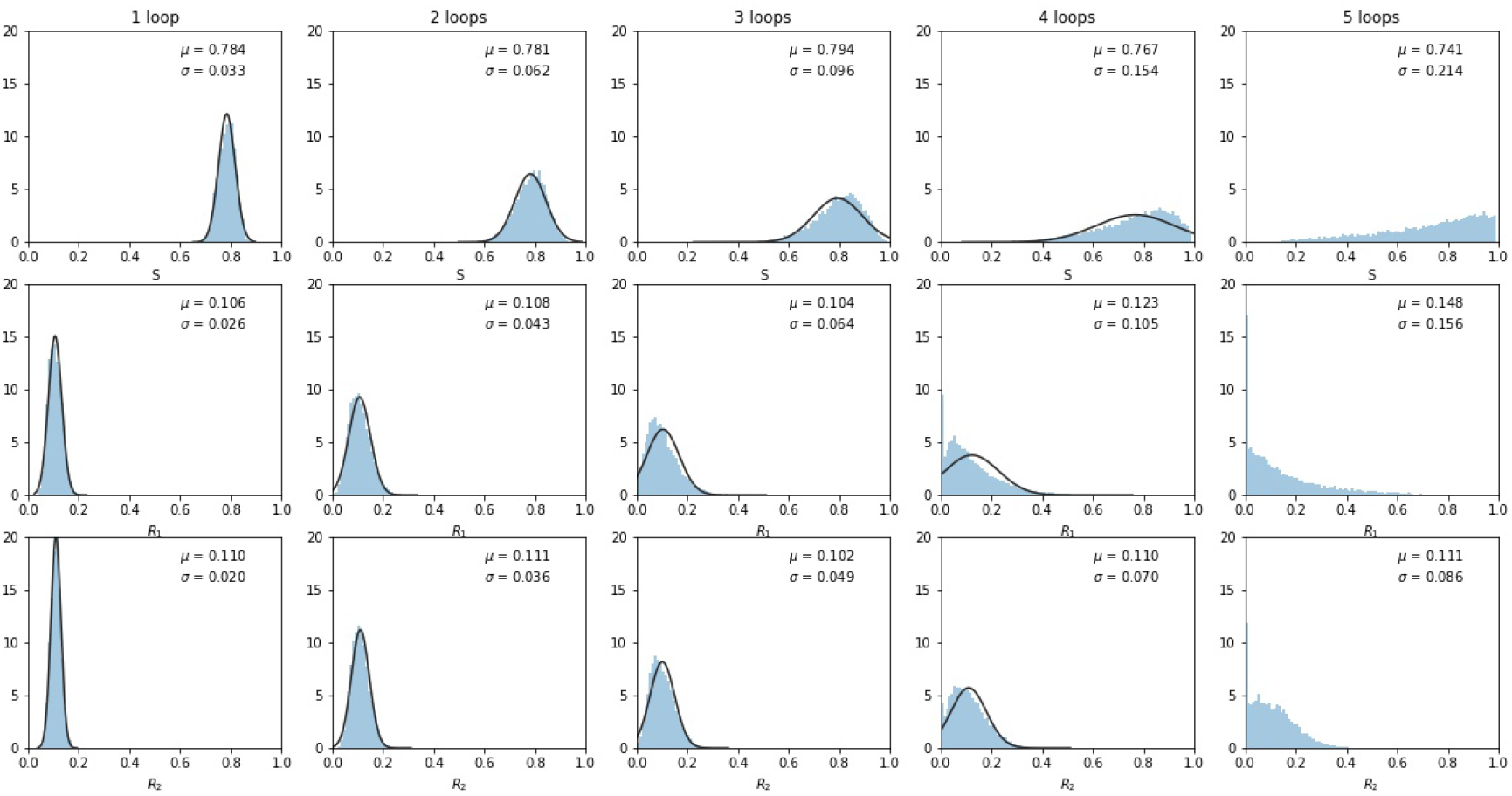
Histogrammed distributions of the three subpopulations as loops increase. Note that some tumors began to fill at the *S* = 1 and *R*_1_ = 1 corners, distorting the multivariate Gaussian nature of the distributions. Mean *µ* and standard deviations *σ* are shown.

The histogrammed distributions for synergistic (*e >* 0) and antagonistic (*e <* 0) drug interactions are shown in figure 6(a),(b). In both cases, the distribution begins to distort after around three loops, with the synergistic interactions breaking down slightly sooner in the third loop. This is shown nicely in figure 7 which depicts the two principal components (on a log-linear plot) as a function of cycle number for additive, synergistic, and antagonistic interactions. In all three cases, the variance of the distribution grows exponentially with loop number, with antagonistic drug interactions outperforming additive interactions, which both outperform synergistic interactions which break down the quickest. One could argue this is because antagonistic interactions are able to suppress competitive release of the resistant populations more efficiently than synergistic interactions, as discussed in [26].

**FIG. 6.**
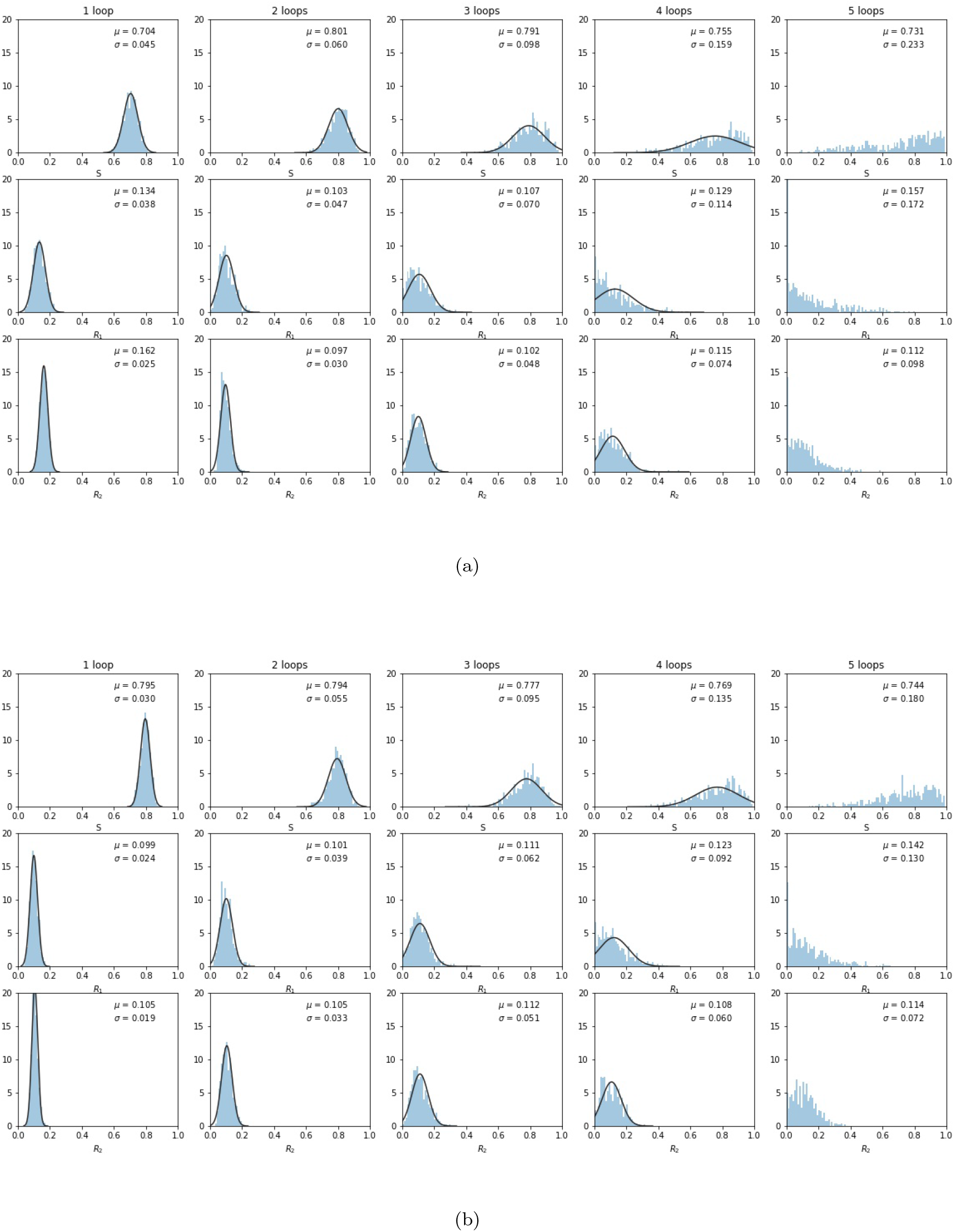
Same as figure 5, but for synergistic *e >* 0 and antagonistic *e <* 0 drug interactions. (a) *e* = 0.3; (b) *e* = −0.3.

**FIG. 7.**
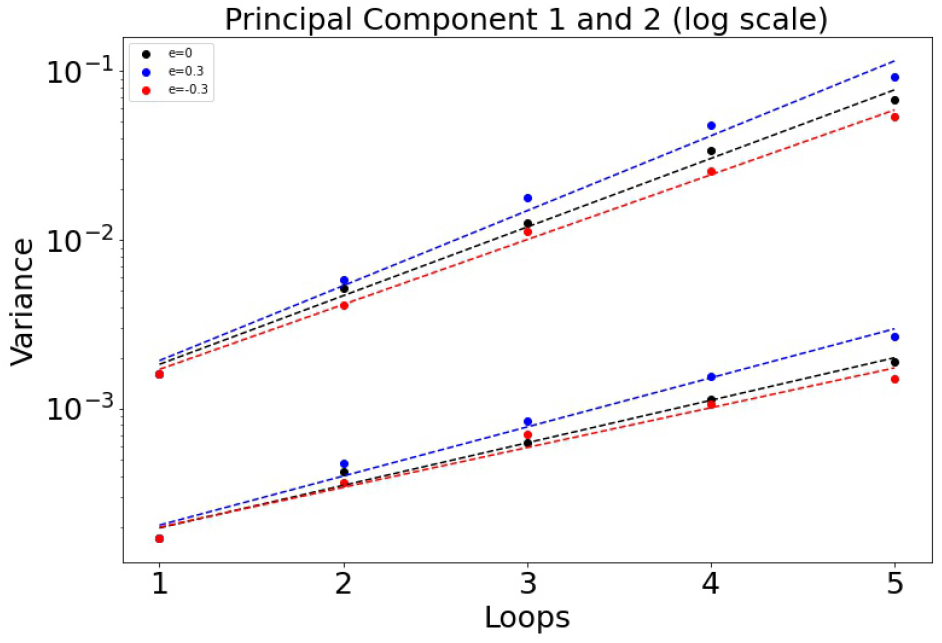
Variance *σ*_*e*_ of the principal components as a function of the cycle number *n* plotted on log-linear axes, so *σ*_*e*_ ∼*a* exp (*α*_*e*_*n*). Note that antagonistic interactions grow the slowest, while synergistic interactions grow fastest. *α*_0_: 0.936 (PCA 1) and 0.580 (PCA 2). *α*_0.3_: 1.02 and 0.670. *α*_*−*0.3_: 0.885 and 0.543.

**FIG. 8.**
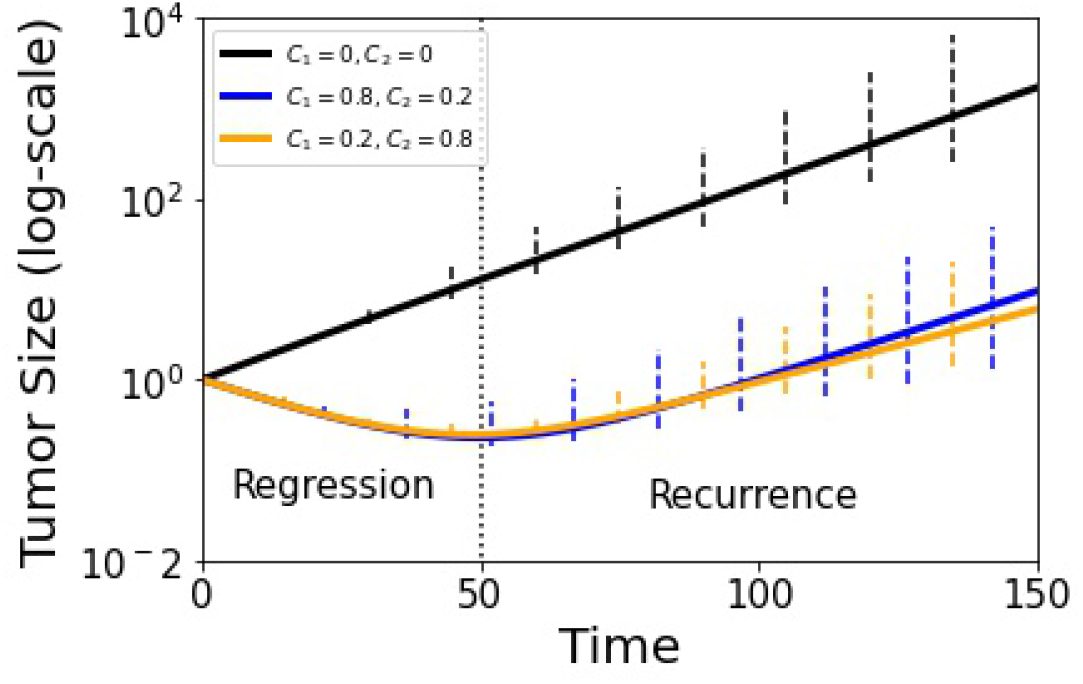
Tumor growth curve with adjusted replicator model using *g* = 1.05 and the stochastic Moran process model (100 runs). Black curve shows untreated growth, blue and yellow show adaptive therapy results. Error bars indicate 90% confidence level. Tumor recurrence sets in at *t* ≈ 50 in dimensionless time units.

## IV. DISCUSSION

The exponential increase in the distribution variance with each adaptive cycle degrades the effectiveness of each subsequent round of chemotherapy since the cycles were designed to stay in a closed evolutionary loop only in the deterministic limit *N*→ ∞. Ideally, the time-dependent chemoschedule keeps the three subpopulations in long-term competition with each other to avoid chemoresistance indefinitely, although delaying chemoresistance through several cycles may be all that is practically achievable due to stochastic fluctuations. Nonetheless, we point to several strategies that could help extend the timeframe over which the schedules could remain effective, even if the evolutionary loops are not closed. Three strategies are discussed in [36]. The first is to explore other possibilities than the commonly used and accepted 50% rule which identifies a nominal baseline tumor size and advocates applying chemotherapy until the size shrinks by half. Then, when the tumor reaches the nominal baseline size, chemotherapy is again implemented for another cycle, etc. It is easy to imagine that there could be better, more optimized options, but none have been clinically tested as far as we are aware. The second is to explore the role of tumor size in deciding when to apply chemotherapy. Typically, chemotherapy commences as soon as a tumor is discovered, but it is conceivable that this may not be optimal in all cases. In [5] we explored using different chemotherapy schedules for different tumor growth rates, but these ideas have not yet been tested in the clinic. The third idea discussed in [36] is to explore the optimal frequency distribution of the cell populations in which to begin adaptive therapy. From the point of view of our mathematical model, these ideas generally relate to the size and location of the *ABCA* cycle in the tri-linear phase plane that would be optimal for an adaptive schedule. We, of course, have discretion over both where in the tri-linear plane the cycle starts (location of point *A* in figure 2(a)), and how large an area the evolutionary loop *ABCA* should enclose, both of which are tied to the suggestions made in [36]. Optimizing these decisions could well have medical implications that could and should be explored through clinical trials.

## ACKNOWLEDGMENTS

We gratefully acknowledge support from the Army Research Office MURI Award #W911NF1910269 (2019-2024).

